## Supplementary Materials for "Antibiotic collateral sensitivity is contingent on the repeatability of evolution"

Nichol et al.

#### Supplementary Note 1

The value of $r$ in the mathematical model determines the extent to which the fitness advantage of a mutation biases the likelihood that it is the next genotype to fix in the population. Throughout our analysis we take $r = 0$, corresponding to a random walk on the fitness landscape wherein the next population genotype is chosen randomly amongst all neighbouring fitter genotypes. This value of $r$ corresponds to the models proposed previously [1, 2, 3].

An alternative model is to take $r = 1$ [4, 5]. Under such a model our results are qualitatively unchanged. Of the 82,944 unique tables of collateral response the most likely occurs with probability 0.0097 for $r = 1$ (0.0023 for $r = 0$). Amongst the 225 ordered drug pairs with $r = 1$ we still find only 28 with guaranteed collateral sensitivity, 94 with guaranteed cross resistance, 15 for which the first drug makes no difference, and 88 for which the first drug can induce either collateral sensitivity or cross resistance in the second. If a collateral response table is generated by stochastic *in silico* simulation, and a collaterally sensitive drug pair chosen at random, then the first of these two drugs will induce cross resistance in the second with expected probability 0.548 (0.513 for $r = 0$) as determined from $10^6$ simulations of this process.

Note that as $r \to \infty$, the biased random walk becomes a deterministic walk in which the fittest neighbouring genotype always achieves fixation. In this case, evolution is always repeatable and the collateral response is stable. The model $r \to \infty$ has been proposed previously for protein evolution [6, 7], but is likely inappropriate for modelling evolution in asexually reproducing populations, as evidenced by our own observations of evolutionary divergence during experimental evolution.

### Supplementary Tables

| Antibiotic | Abbreviation | Group | Concentration |
|---|---|---|---|
| Ampicillin | AMP | Aminopenicillin | 2,048 $\mu$g ml$^{-1}$ [8] |
| Amoxicillin | AM | Aminopenicillin | 512 $\mu$g ml$^{-1}$ |
| Cefaclor | CEC | Cephalosporin | 1 $\mu$g ml$^{-1}$ |
| Cefotaxime | CTX | Cephalosporin | 0.05 $\mu$g ml$^{-1}$ |
| Ceftizoxime | ZOX | Cephalosporin | 0.03 $\mu$g ml$^{-1}$ |
| Cefuroxime | CXM | Cephalosporin | 1.5 $\mu$g ml$^{-1}$ |
| Ceftriaxone | CRO | Cephalosporin | 0.045 $\mu$g ml$^{-1}$ |
| Amoxicillin + Clavulanic acid | AMC | Penicillin derivative + $\beta$-Lactamase inhibitor | 512 $\mu$g ml$^{-1}$ (Amoxicillin) and 8 $\mu$g ml$^{-1}$ (Clav) |
| Ceftazidime | CAZ | Cephalosporin | 0.1 $\mu$g ml$^{-1}$ |
| Cefotetan | CTT | Cephalosporin | 0.312 $\mu$g ml$^{-1}$ |
| Ampicillin + Sulbactam | SAM | Penicillin derivative + $\beta$-Lactamase inhibitor | 8 $\mu$g ml$^{-1}$ (Ampicillin) and 8 $\mu$g ml$^{-1}$ (Sulbactam) |
| Cefprozil | CPR | Cephalosporin | 100 $\mu$g ml$^{-1}$ |
| Cefpodoxime | CPD | Cephalosporin | 2 $\mu$g ml$^{-1}$ |
| Pipercillin + Tazobactam | TZP | Penicillin derivative + $\beta$-Lactamase inhibitor | 12 $\mu$g ml$^{-1}$ (Pipercillin) and 8 $\mu$g ml$^{-1}$ (Tazobactam) [8] |
| Cefepime | FEP | Cephalosporin | 0.0156 $\mu$g ml$^{-1}$ |

**Supplementary Table 1:** 15 $\beta$–lactam antibiotics for which Mira et al. [9] derived fitness landscapes.

### Supplementary Figures

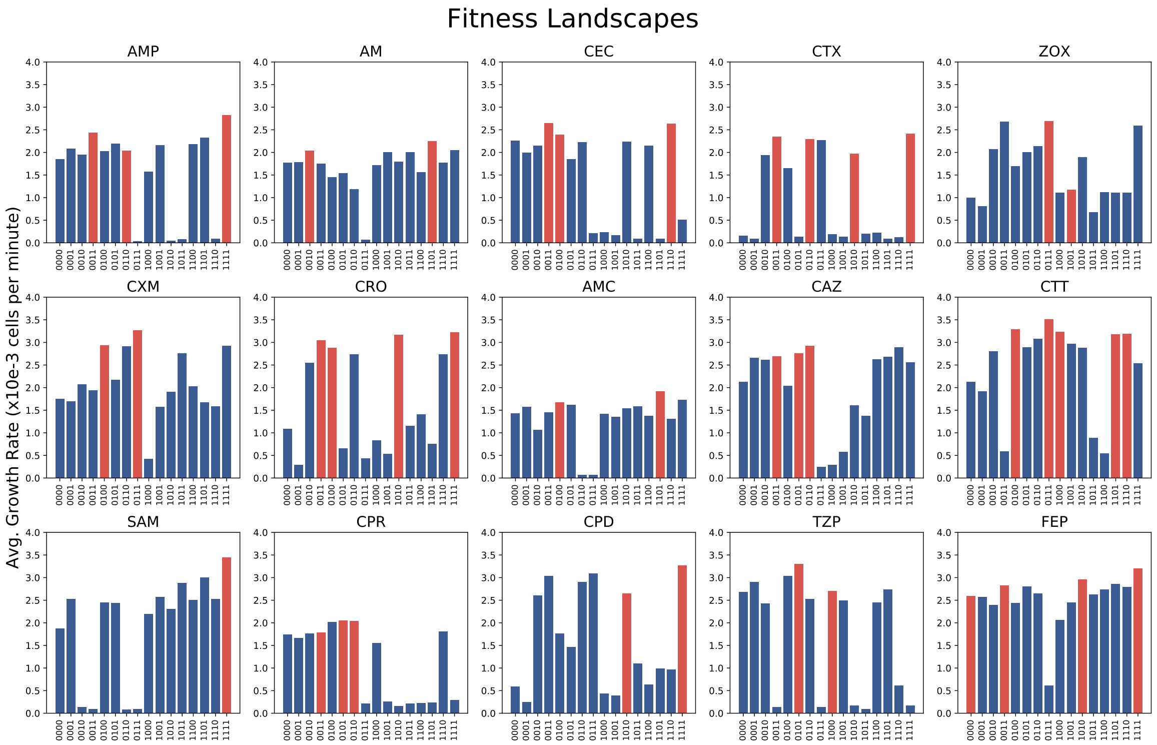


**Supplementary Figure 1:** Fitness landscapes for the 15 antibiotics used in the mathematical model. Red bars indicate fitness peaks (note that adjacent bars need not be adjacent genotypes). These landscapes were originally derived by Mira et al. [9].

Below we present figures showing the the full extent of non–repeatable evolution for the 15 drugs. Each figure corresponds to five first–line and five second–line drugs and consists of 25 subplots, one for each pair. Points in these subplots represent accessible local optima genotypes in the fitness landscape indicated on the $x$-axis. Each point has an $x$ value corresponding to the likelihood that it arises from evolution (from $g_0 = 0000$) in the landscape of the labelled drug, as determined by the mathematical model (with $r = 0$, see Materials and Methods for details). The $y$ value in each subplot corresponds to the fitness of that genotype in a second landscape (as labelled). The wild–type ($g = 0000$) fitness (solid line) and expected fitness following evolution under the first drug (dashed) line are shown. Lines and points are coloured according to the fold–change from wild–type fitness (colour bar).

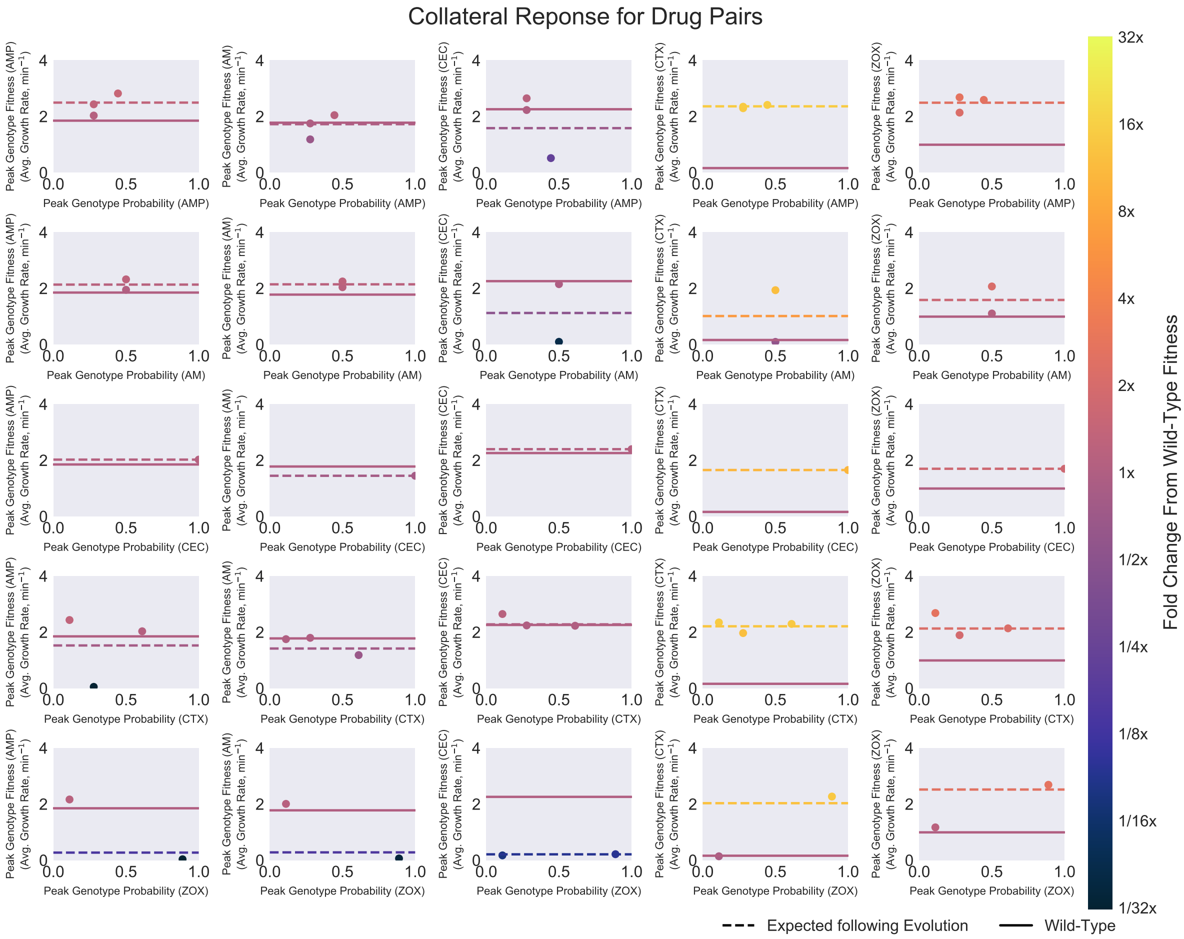


**Supplementary Figure 2: Partial representation of all collateral response possibilities.** Each point in a subplot denotes a peak genotype of the landscape named in the $x$-axis. The $x$-value is the likelihood of evolution proceeding to this peak genotype from $g = 0000$ ($r = 0$ model) under that landscape. The $y$-value is fitness of the peak genotype in the second landscapes named on the $y$-axis. Points are coloured by the fold change sensitivity (colour bar). The solid line delineates the fitness of $g = 0000$ in the second landscape, the dashed line delineates the average fitness following evolution under the first landscape.

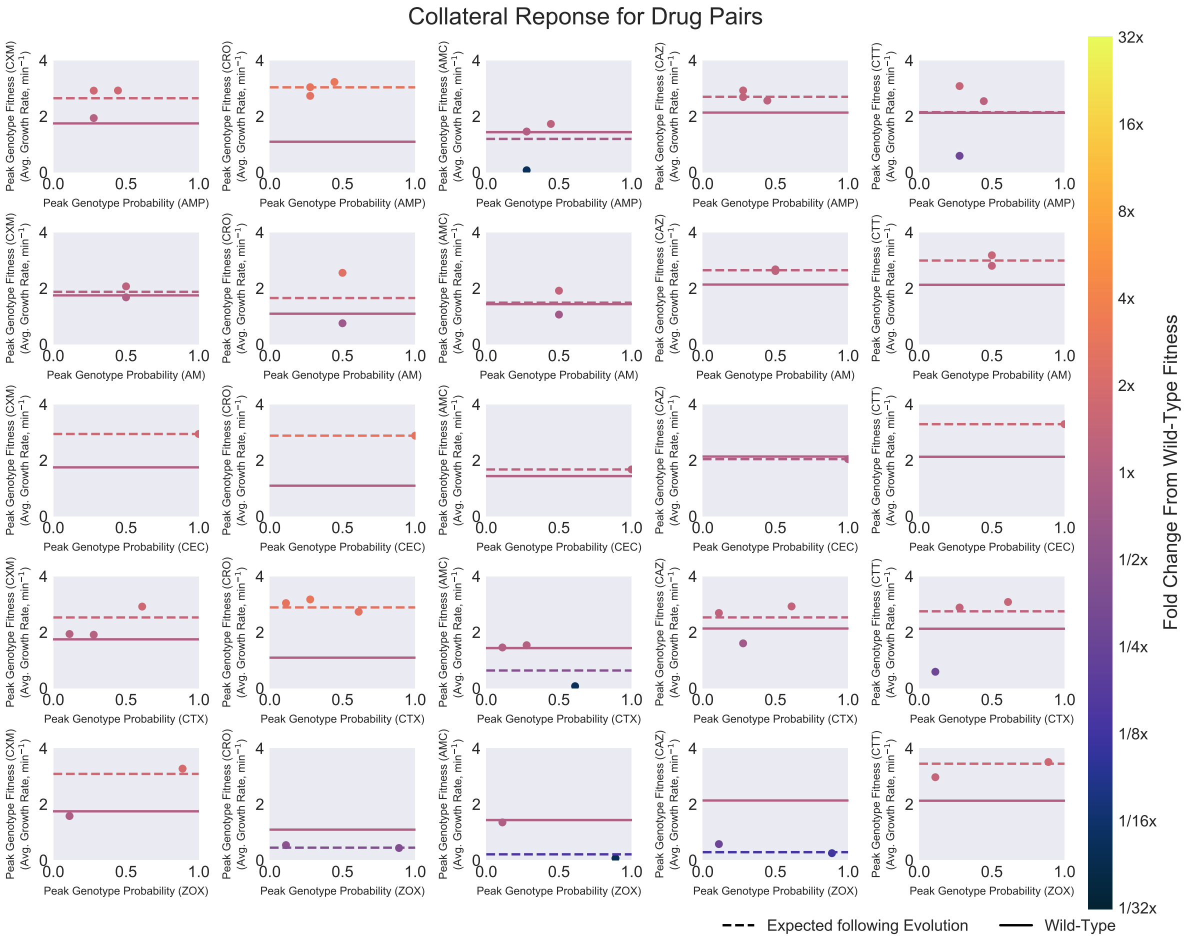


**Supplementary Figure 3: Partial representation of all collateral response possibilities.** Each point in a subplot denotes a peak genotype of the landscape named in the $x$-axis. The $x$-value is the likelihood of evolution proceeding to this peak genotype from $g = 0000$ ($r = 0$ model) under that landscape. The $y$-value is fitness of the peak genotype in the second landscapes named on the $y$-axis. Points are coloured by the fold change sensitivity (colour bar). The solid line delineates the fitness of $g = 0000$ in the second landscape, the dashed line delineates the average fitness following evolution under the first landscape.

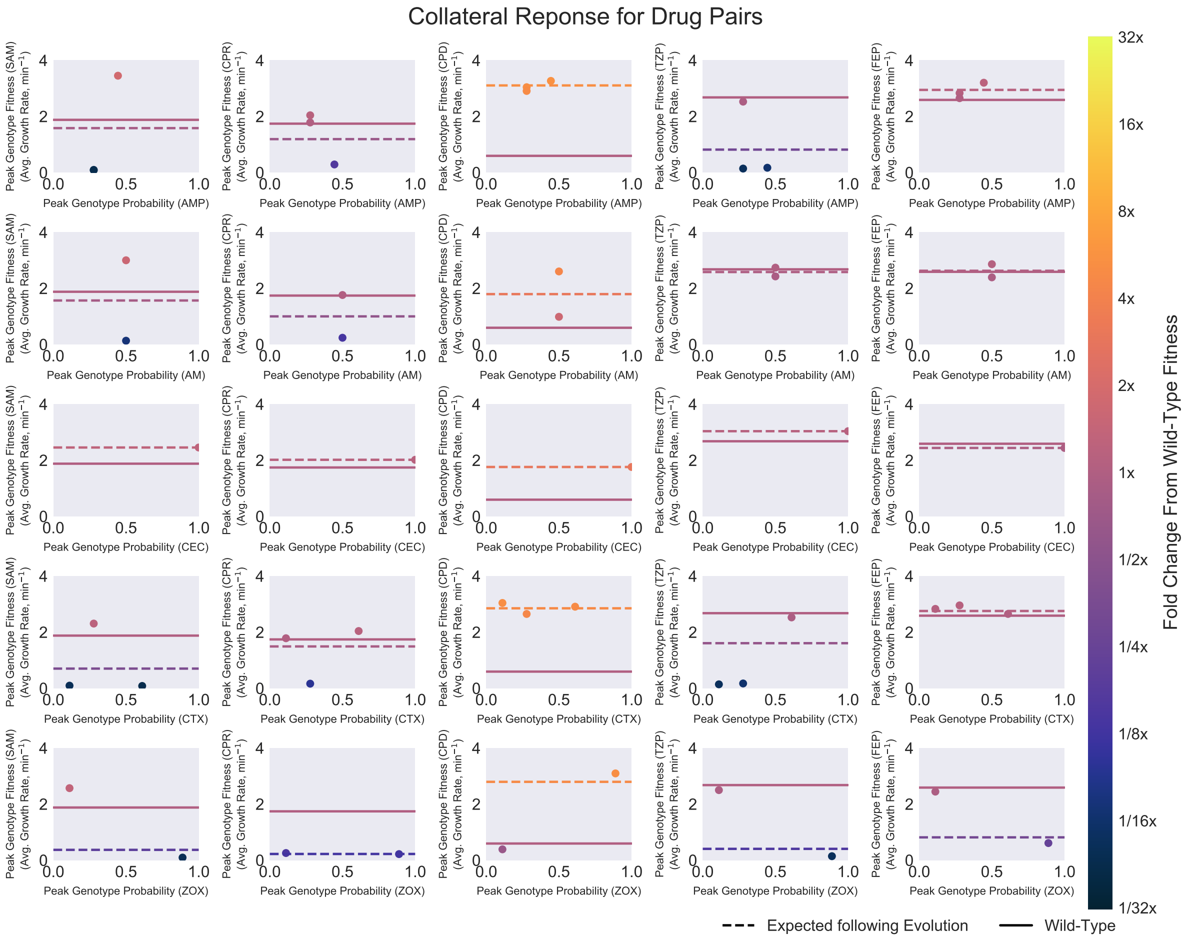


**Supplementary Figure 4: Partial representation of all collateral response possibilities.** Each point in a subplot denotes a peak genotype of the landscape named in the $x$-axis. The $x$-value is the likelihood of evolution proceeding to this peak genotype from $g = 0000$ ($r = 0$ model) under that landscape. The $y$-value is fitness of the peak genotype in the second landscapes named on the $y$-axis. Points are coloured by the fold change sensitivity (colour bar). The solid line delineates the fitness of $g = 0000$ in the second landscape, the dashed line delineates the average fitness following evolution under the first landscape.

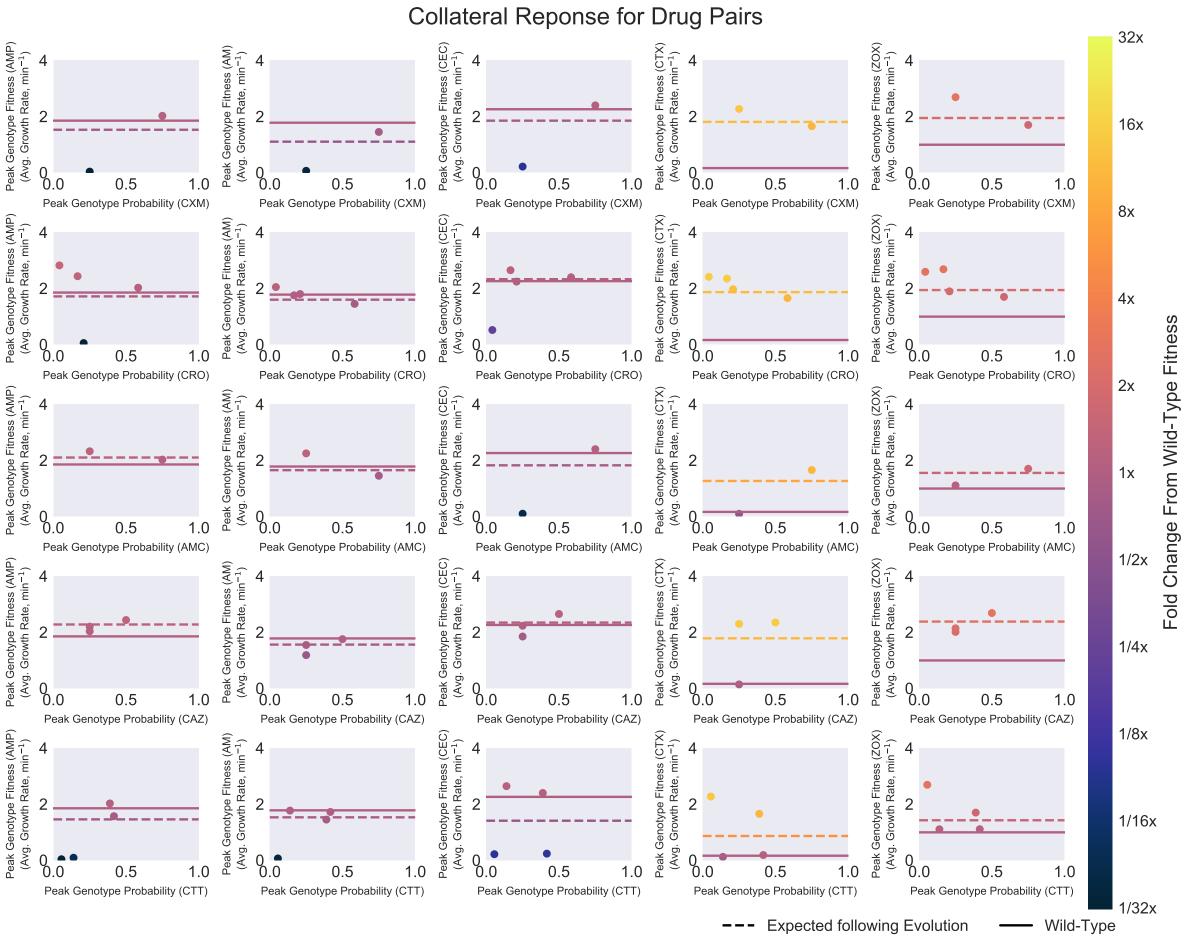


**Supplementary Figure 5: Partial representation of all collateral response possibilities.** Each point in a subplot denotes a peak genotype of the landscape named in the $x$-axis. The $x$-value is the likelihood of evolution proceeding to this peak genotype from $g = 0000$ ($r = 0$ model) under that landscape. The $y$-value is fitness of the peak genotype in the second landscapes named on the $y$-axis. Points are coloured by the fold change sensitivity (colour bar). The solid line delineates the fitness of $g = 0000$ in the second landscape, the dashed line delineates the average fitness following evolution under the first landscape.

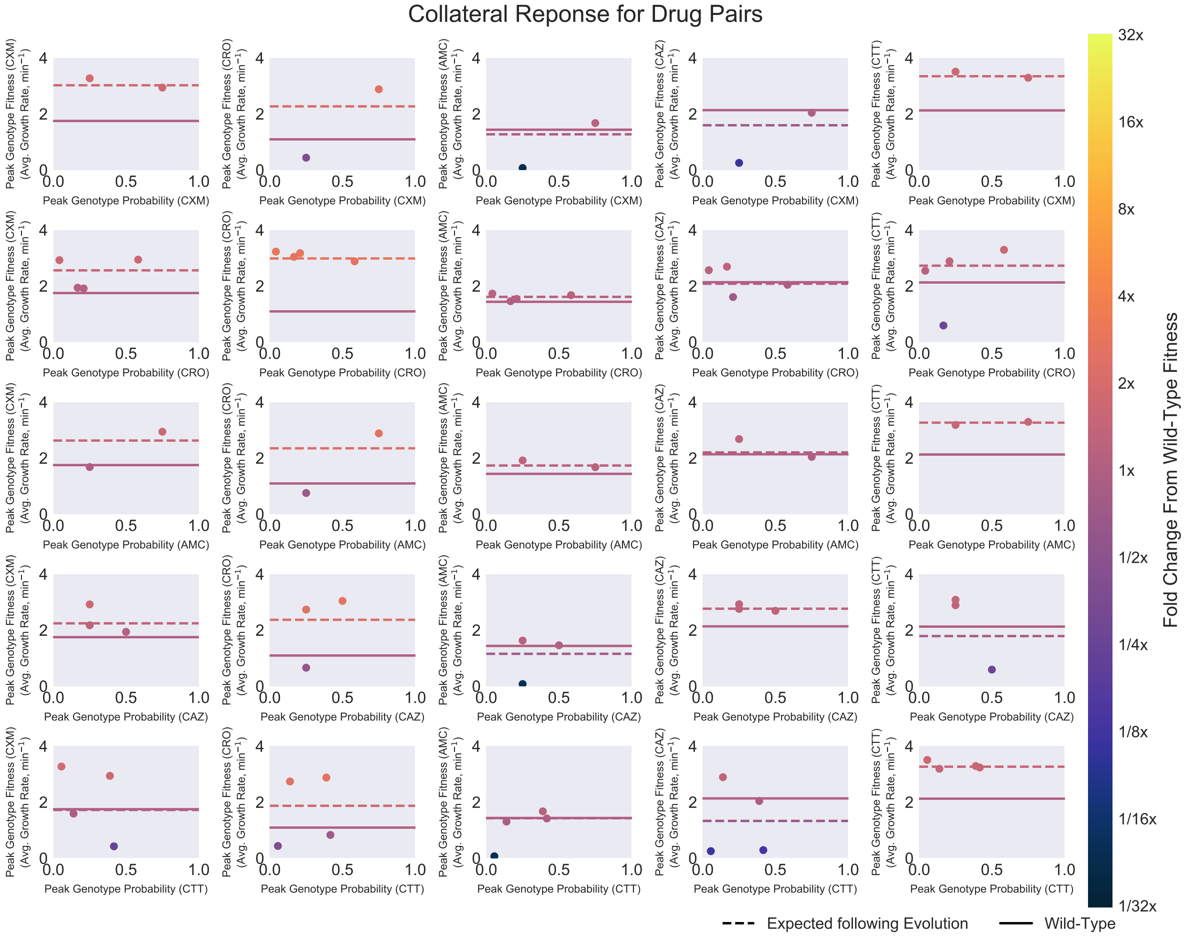


**Supplementary Figure 6: Partial representation of all collateral response possibilities.** Each point in a subplot denotes a peak genotype of the landscape named in the $x$-axis. The $x$-value is the likelihood of evolution proceeding to this peak genotype from $g = 0000$ ($r = 0$ model) under that landscape. The $y$-value is fitness of the peak genotype in the second landscapes named on the $y$-axis. Points are coloured by the fold change sensitivity (colour bar). The solid line delineates the fitness of $g = 0000$ in the second landscape, the dashed line delineates the average fitness following evolution under the first landscape.

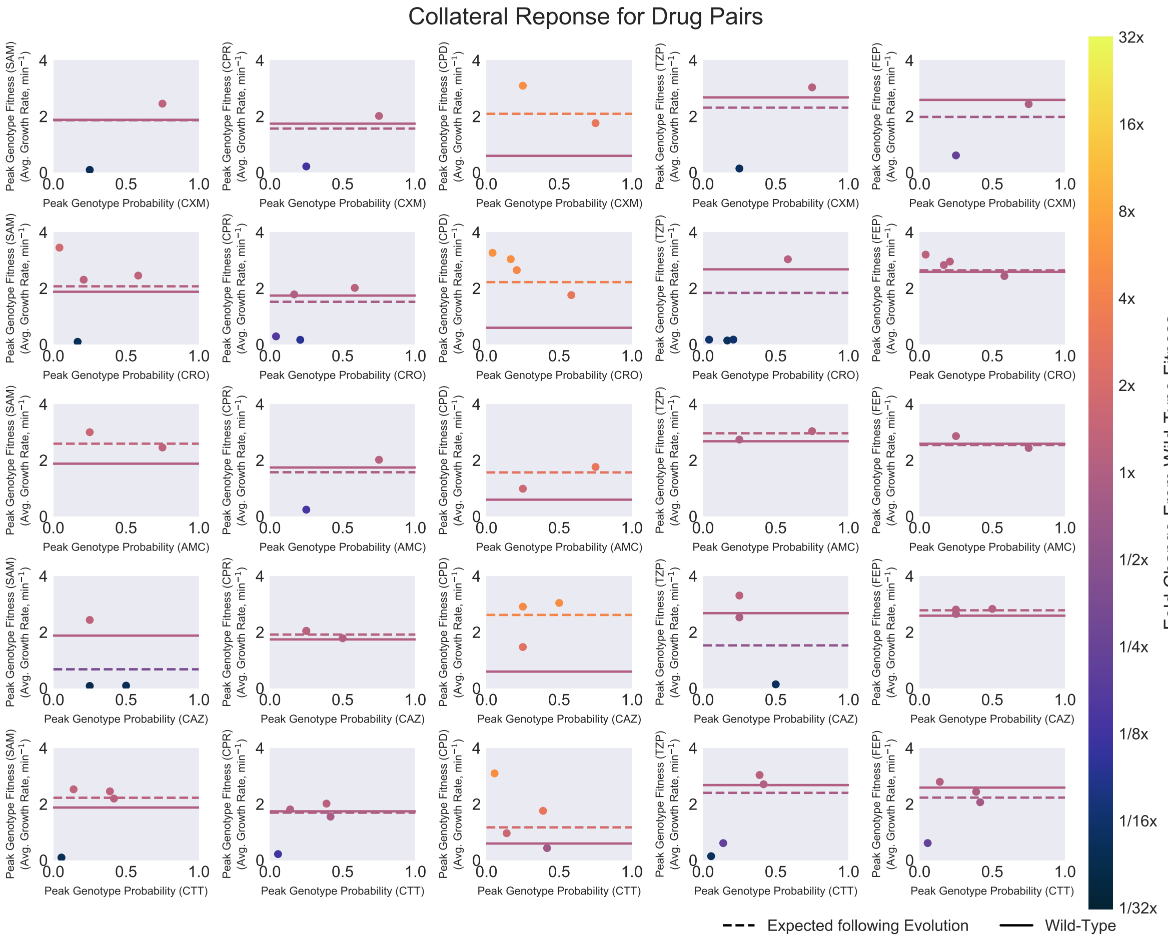


**Supplementary Figure 7: Partial representation of all collateral response possibilities.**Each point in a subplot denotes a peak genotype of the landscape named in the $x$-axis. The $x$-value is the likelihood of evolution proceeding to this peak genotype from $g = 0000$ ($r = 0$ model) under that landscape. The $y$-value is fitness of the peak genotype in the second landscapes named on the $y$-axis. Points are coloured by the fold change sensitivity (colour bar). The solid line delineates the fitness of $g = 0000$ in the second landscape, the dashed line delineates the average fitness following evolution under the first landscape.

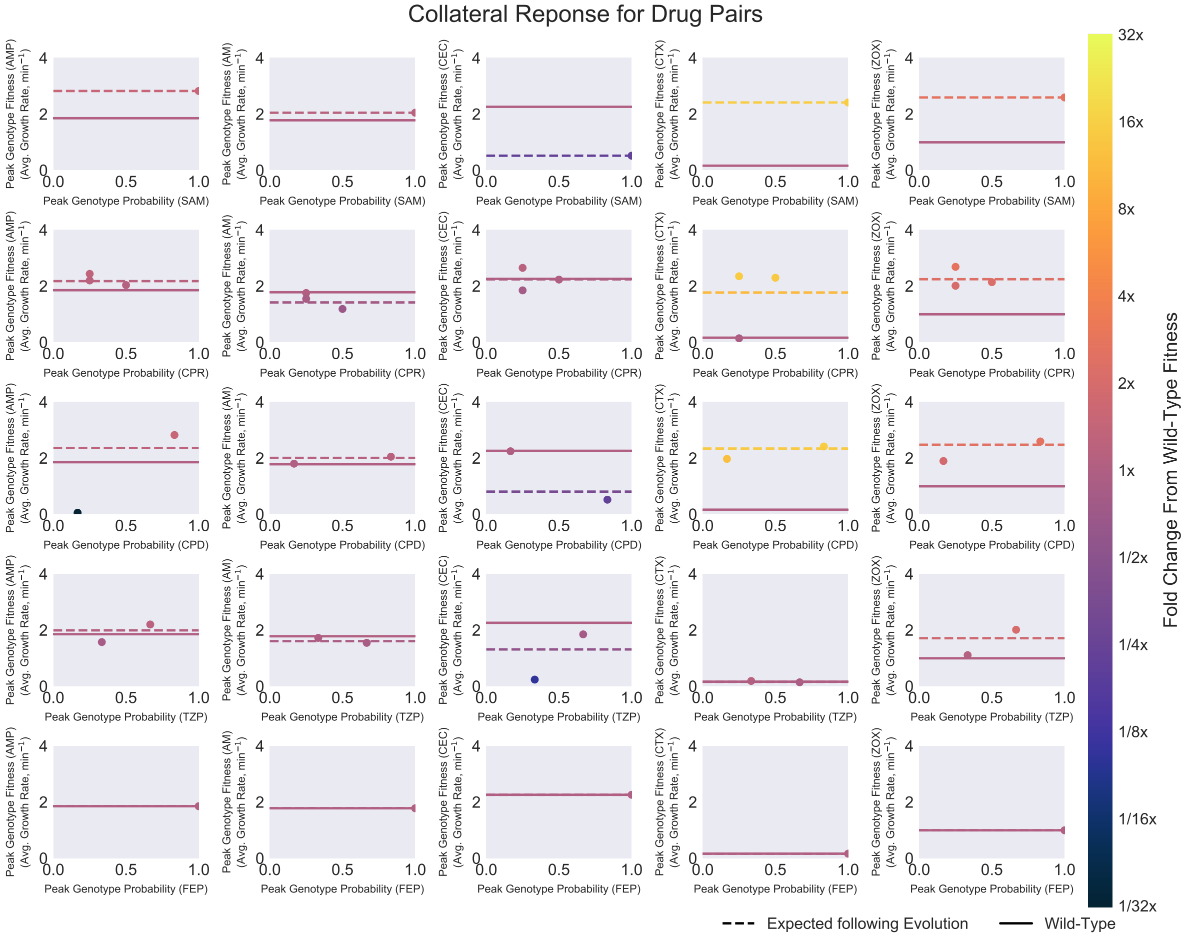


**Supplementary Figure 8: Partial representation of all collateral response possibilities.** Each point in a subplot denotes a peak genotype of the landscape named in the $x$-axis. The $x$-value is the likelihood of evolution proceeding to this peak genotype from $g = 0000$ ($r = 0$ model) under that landscape. The $y$-value is fitness of the peak genotype in the second landscapes named on the $y$-axis. Points are coloured by the fold change sensitivity (colour bar). The solid line delineates the fitness of $g = 0000$ in the second landscape, the dashed line delineates the average fitness following evolution under the first landscape.

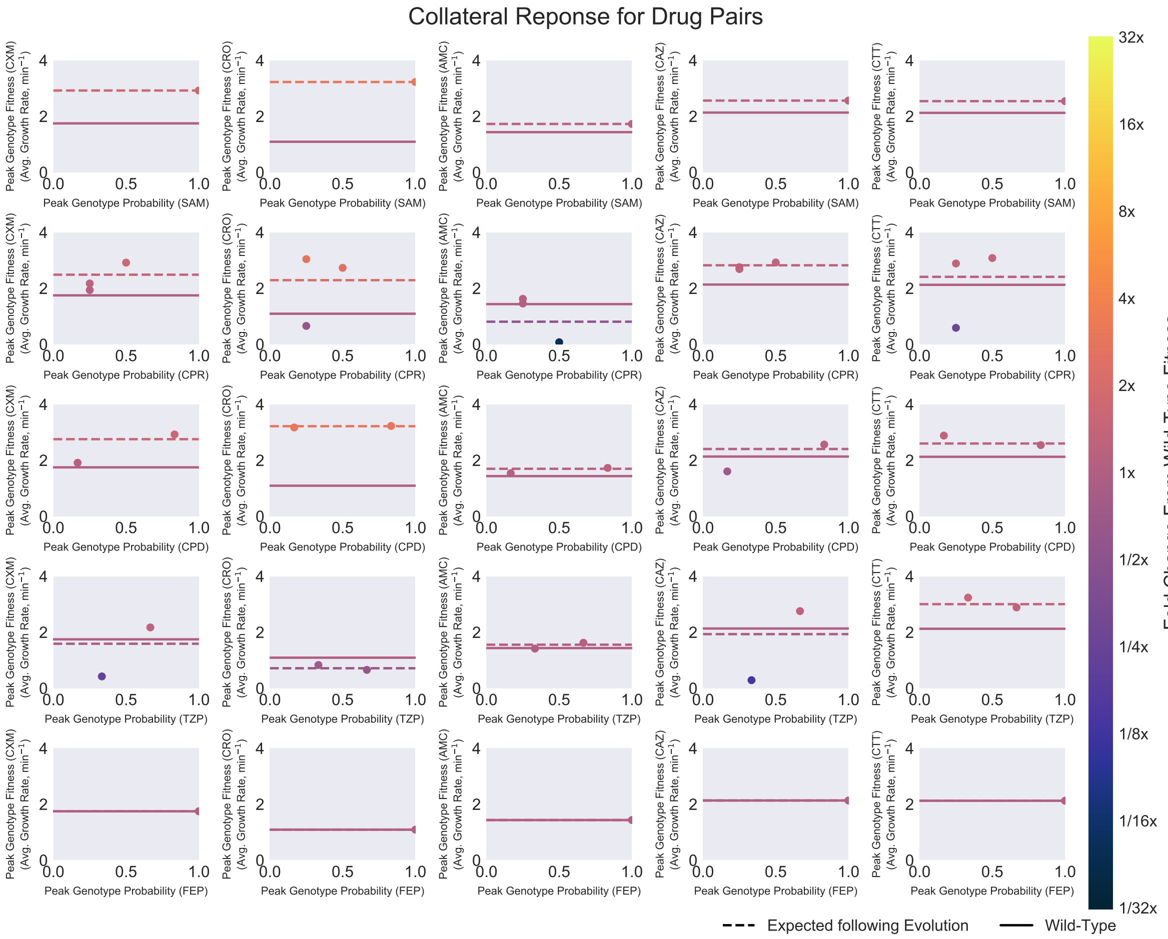


**Supplementary Figure 9: Partial representation of all collateral response possibilities.** Each point in a subplot denotes a peak genotype of the landscape named in the $x$-axis. The $x$-value is the likelihood of evolution proceeding to this peak genotype from $g = 0000$ ($r = 0$ model) under that landscape. The $y$-value is fitness of the peak genotype in the second landscapes named on the $y$-axis. Points are coloured by the fold change sensitivity (colour bar). The solid line delineates the fitness of $g = 0000$ in the second landscape, the dashed line delineates the average fitness following evolution under the first landscape.

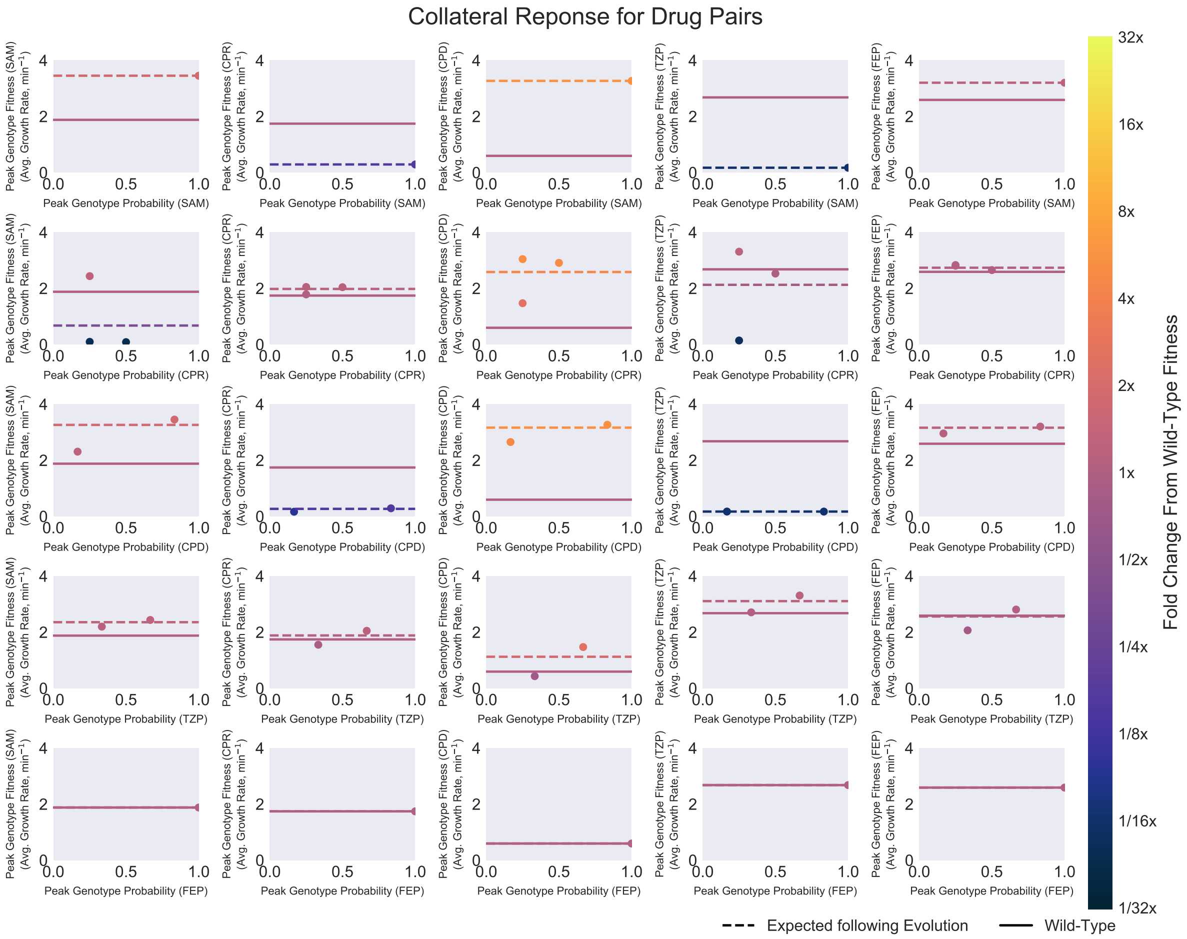


**Supplementary Figure 10: Partial representation of all collateral response possibilities.** Each point in a subplot denotes a peak genotype of the landscape named in the $x$-axis. The $x$-value is the likelihood of evolution proceeding to this peak genotype from $g = 0000$ ($r = 0$ model) under that landscape. The $y$-value is fitness of the peak genotype in the second landscapes named on the $y$-axis. Points are coloured by the fold change sensitivity (colour bar). The solid line delineates the fitness of $g = 0000$ in the second landscape, the dashed line delineates the average fitness following evolution under the first landscape.

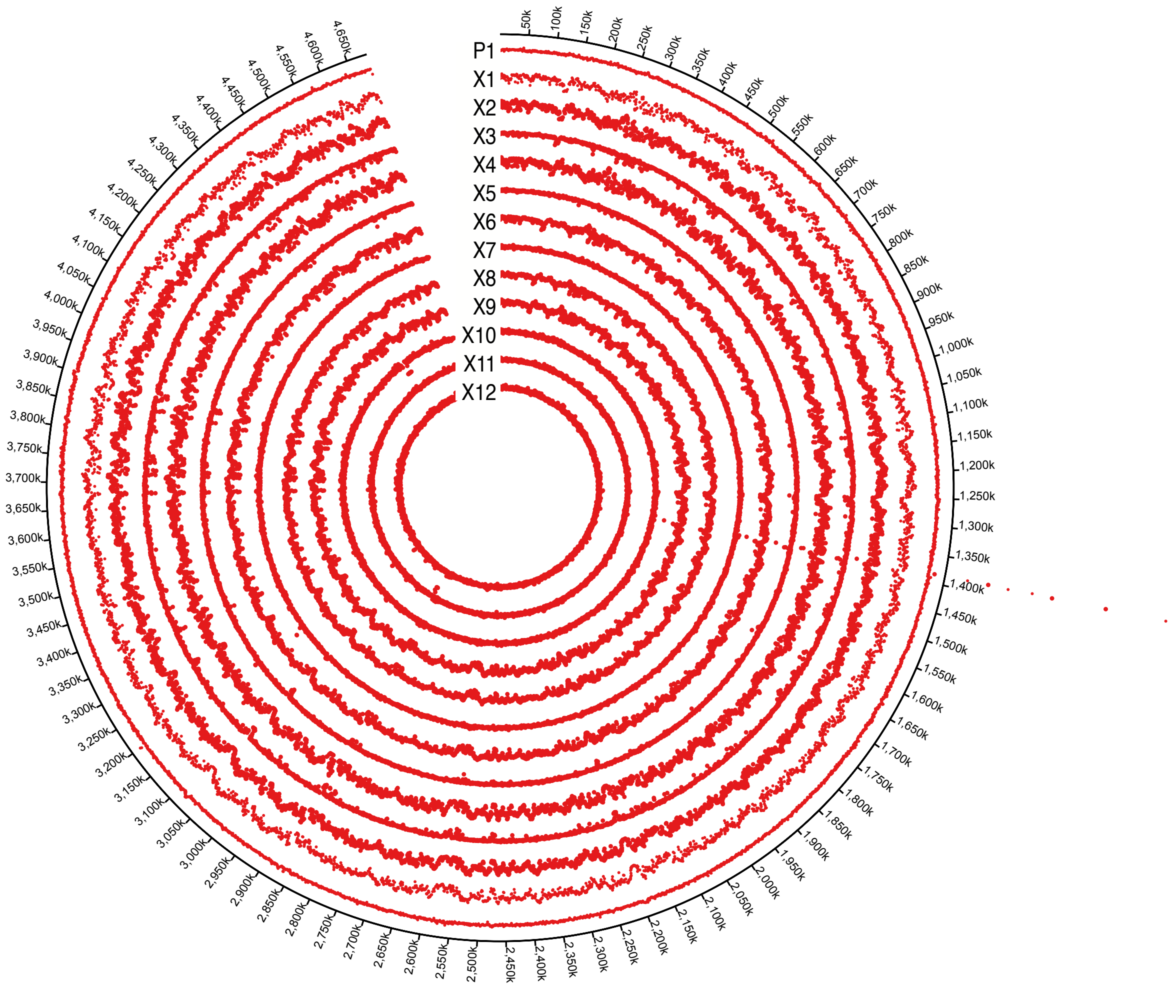


**Supplementary Figure 11:** Circos plots show the aligned read depths for the whole genome sequencing of P1 and X1-X12
